# Reaction-Diffusion Model of PDGF-driven Recruitment in Experimental Glioblastoma

**DOI:** 10.1101/038471

**Authors:** Susan Christine Massey, Kristin R. Swanson

## Abstract

Platelet-derived growth factor (PDGF) drives the formation of gliomas in an experimental animal model, which notably involves the recruitment of large numbers of glial progenitor cells. (Assanah 2006). In order to understand the underlying mechanism, particularly what factors influence the degree of recruitment, and how varied amounts of PDGF would affect the gross characteristics and overall appearance of tumors in the brain, we adapted a reaction diffusion model of glioma, which has been used for analyzing clinical data, to model the interactions at play in these experimental models.

## I. PLATELET-DERIVED GROWTH FACTOR MEDIATED RECRUITMENT IN EXPERIMENTAL GLIOBLASTOMA

To elucidate platelet-derived growth factor mediated tumor growth, a retrovirus expressing platelet-derived growth factor B homodimer (PDGF-BB) and green fluorescent protein (GFP) was injected into rat brains. Brain slices of sacrificed animals were incubated and imaged via fluorescence microscopy. Microscopy revealed that the virus transfected oligodendrocyte progenitor cells.

Interestingly, after 17 days, tumors were composed of both transfected and normal glial progenitor cells (the normal glial progenitors only expressing the red cherry, having no GFP to indicate transfection). Surprisingly, tumors were 80% un-transfected glial progenitor cells, suggesting that the increased secretion of PDGF-BB by the transfected cells “recruited” un-transfected progenitors via paracrine signaling, raising their proliferation and migration rates such that they behaved as cancer cells. Moreover, the tumors look morphologically similar to human glioblastoma (GBM), which is diffusely infiltrative at the outer margins [2]-[4].

Following these results, we wondered about the possible role of recruitment in human GBM - were the growth rates of these experimental tumors too fast to correspond to observations in the clinical setting? And how would the characteristics of a tumor change if the degree of recruitment was different?

## II. ILLUSTRATIVE RESULTS OF APPLICATION OF METHODS

Using a reaction-diffusion model, we were able to show that a simple recruitment mechanism could substantially explain the experimental observations, achieving 80% recruitment and the same size at 17 days post injection with virus (Fig. 1 A). Furthermore, we found that the level of PDGF production/secretion affects the invasive profile of the tumor, as well as the rate of radial growth, following the path of outward expansion of a given cellular density (Fig. 1 B). Growth rates (Fig. 1 C) correspond well to the clinical human data, as well. These results and others are published in [5].

**Figure 1.**
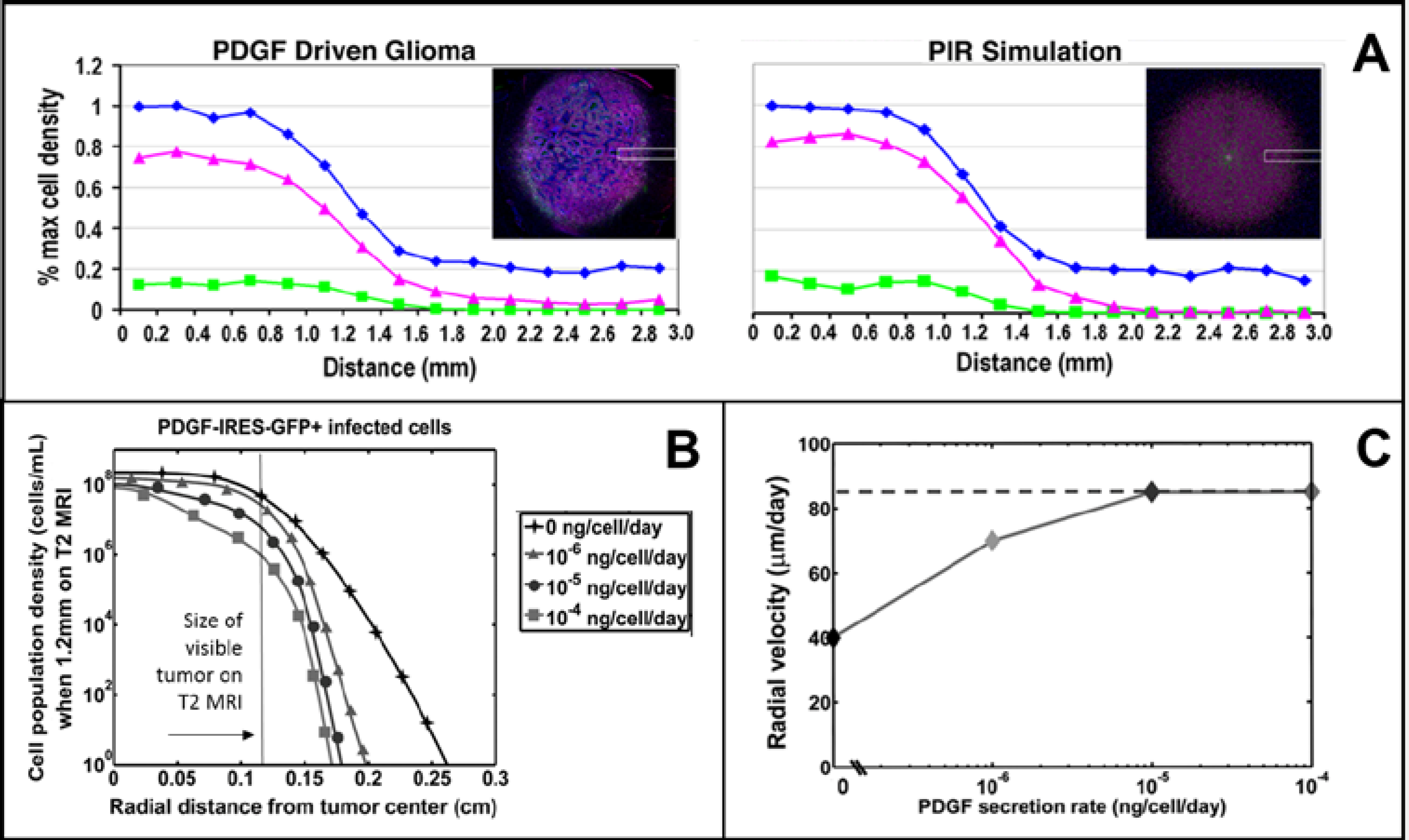
Fluorescence microscopy and simulation. Visualizing simulation results to match fluorescence microscopy images as well as plotting the cell count data (here as a percent of maximum cell density) shows good agreement between the mathematical model and the experimental data. The blue line represents DAPI labeled cells (all cells), the magenta line — cells co-labeled with DAAPI and olig2+, and the green line — GFP labeled transfected cells. B. Cell population densities for varied degrees of PDGF secretion. Legend shows line markers that correspond to rates of PDGF secretion for the simulation results. Note that at a fixed size on MRI, the simulated tumors with no PDGF secretion (exclusively autocrine stimulated) extend further beyond the MR-visible region than any other. C. Radial velocity of simulated tumors, at varied levels of PDGF secretion. For no paracrine PDGF signaling (0 ng/cell/day secreted), the radial growth velocity is approx. 40 um/day. This radial growth rate increases for higher levels of signaling, but levels off at a max rate. All of these tumor radial growth velocities fall within the range of those observed in human GBMs.

## III. QUICK GUIDE TO REACTION DIFFUSION METHODS

Our initial objective was to confirm that this recruitment mechanism could explain the experimental observations in [1], and secondarily, to try to understand what factors influence this process and the degree of recruitment. To do this, we adapted the proliferation-invasion (PI) model developed by Swanson et al [2], [3], [6], [7]. This is a reaction diffusion model, which is useful for understanding the growth of tumors at the tissue/organ level [8]-[10], making it suitable for clinical applications [11]-[15], as well.

### A. Reaction Diffusion Models

A reaction diffusion model is a partial differential equation, with variables that vary in both time and space. These models follow the general pattern where the time derivative of the variable is set equal to the sum of two types of terms: 1) *diffusion terms:* terms involving the second spatial derivative of the variable; and 2) *reaction terms:* terms involving the variables with no derivatives. Both kinds of terms may or may not have coefficients. The PI model uses the reaction term to describe the cellular proliferation component of tumor, and the diffusion term models the invasive migration component of tumor growth. This has been particularly useful for modeling gliomas, whose infiltration resembles a diffusion process when cell tracks are analyzed via windrose plots [5].

Since we had more than one population of cells we were interested in, we had two reaction diffusion equations for each of the two cell types relevant to the recruitment story: transfected (Eqn. 1) and recruited (untransfected) glial progenitor cells (Eqn. 2). Furthermore, we determined that it was necessary to explicitly model the PDGF, which mediates the relationship between these two populations. This equation (3) is also a reaction diffusion equation, with reaction terms that account for the production (by transfected cells) and consumption (by both recruited and transfected cells), and the diffusion term to model the movement of the growth factor as it moves into the tissue after being secreted by the transfected cells. Notice that in the equations, the diffusion term is just right of the equality sign, and the remaining terms further right of that are reaction terms. We also accounted for dose response effects to PDGF using Michaelis-Menten terms in the functional form coefficients (see [5] and its associated supplemental material for details).

#### Model Equations

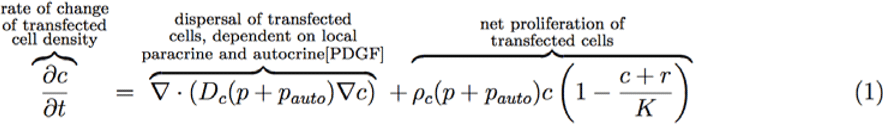

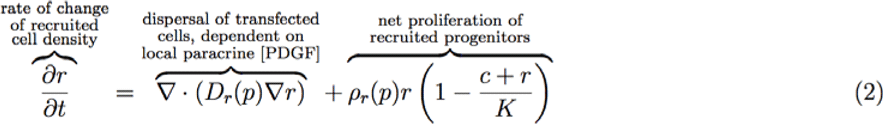

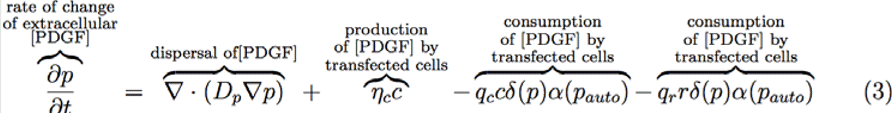

### B. Solving Reaction Diffusion Equations

To solve our model equations, we relied on a numerical method encapsulated in Matlab’s partial differential equation (PDE) solver, the pdepe.m function. This is a useful tool for the beginner, since it has the advantage of selecting the most appropriate numerical method for solving the problem, and it handles reaction-diffusion problems well. To use the solver, we must specify the equations, as well as initial and boundary conditions. These conditions are necessary for solving initial boundary value problems such as our present reaction diffusion model. Using data from the initiation of tumors for the initial condition, we had the initial volume of cells that could be exposed to virus, and assumed a high rate of transfection of the local glial progenitors in that value. For the spatial domain, we used spherical symmetry and allowed the tumor to grow out to 1 cm, the approximate radius of a rat brain inside the skull, at which was a no-flux boundary. We then allowed our simulations to run to 20 days post infection, since the rats with experimental tumors reached morbidity by that time.

### C. Model Parameterization

Many of our parameter coefficients were determined using literature values, but several were not available in literature sources. In the case of parameters with no literature values or direct experimental data available, we explored a range of possibilities and selected the parameter value that resulted in the best model fit of the experimental data. For example, the rate of PDGF secretion by transfected cells, η_c_, is unknown, and we thus ran simulations exploring a range of values over several orders of magnitude to cover the range (and perhaps beyond) of what is physiologically possible. We then compared the results of these simulations with data, and selected the value for η_c_ that corresponded to the model simulation result that best matched the recruitment levels and other experimental data we observed. As demonstrated in the previous section and in [5], testing different values for these unknown parameters led to interesting insights about how these experimental tumors respond to PDGF.

## ACKNOWLEDGMENT

We thank the organizers of the Maths of the ICBPs and PSOCs meeting that inspired the creation of this handbook.

